# Reconstructing tumor trajectories during therapy through integration of multiple measurement modalities

**DOI:** 10.1101/2021.01.14.426737

**Authors:** Jason I. Griffiths, Jinfeng Chen, Onalisa Winblad, Anne O’Dea, Priyanka Sharma, Meghna Trivedi, Kevin Kalinsky, Kari B. Wisinski, Ruth O’Regan, Issam Makhoul, Yuan Yuan, Laura M. Spring, Aditya Bardia, Mohammad Jahanzeb, Frederick R. Adler, Adam L. Cohen, Andrea H. Bild, Qamar J. Khan

## Abstract

**Background:** Accurately determining changes in tumor size during therapy is essential to evaluating response or progression. However, individual imaging methodologies often poorly reflect pathologic response and long-term treatment efficacy in patients with estrogen receptor positive (ER+) early-stage breast cancer. Mathematical models that measure tumor progression over time by integrating diverse imaging and tumor measurement modalities are not currently used but could increase accuracy in measuring response and provide biological insights into cancer evolution.

**Methods:** For ER+ breast cancer patients enrolled on a neoadjuvant clinical trial, we reconstructed their tumor size trajectories during therapy by combining all available information on tumor size, including different imaging modalities, physical examinations and pathological assessment data. Tumor trajectories during six months of treatment were generated, using a Gaussian process and the most probable trajectories were evaluated, based on clinical data, using measurement models that account for biases and differences in precision between tumor measurement methods, such as MRI, ultrasound and mammograms.

**Results:** Reconstruction of tumor trajectories during treatment identified five distinct patterns of tumor size changes, including rebound growth not evident from any single modality. These results increase specificity to distinguish innate or acquired resistance compared to using any single measurement alone. The speed of therapeutic response and extent of subsequent rebound tumor growth quantify sensitivity or resistance in this patient population.

**Conclusions:** Tumor trajectory reconstruction integrating multiple modalities of tumor measurement accurately describes tumor progression on therapy and reveals various patterns of patient responses. Mathematical models can integrate diverse response assessments and account for biases in tumor measurement, thereby providing insights into the timing and rate at which resistance emerges.

## Introduction

Cancer patients’ response to a therapy is highly variable, with some tumors responding slowly or rapidly, others progressing on therapy, and some exhibiting rapid early response followed by rebound regrowth (1, 2). Despite this known diversity of patient tumor trajectories (3, 4) the treatment response of most solid tumors during trials is evaluated by comparing baseline and end point measurements. Patients are then grouped into a small number of response categories based on the best response. Therefore, a tumor that rapidly shrinks by 50% and then rapidly grows back and another tumor that slowly shrinks by 50% are both called partial responses (PR) (5-7). This end-point focused approach overlooks the diversity of tumor responses during a given trial, including what happens in between time points, and fails to distinguish different outcomes such as patients with stable disease versus those with an initial response followed by subsequent disease rebound (8). Due to different intrinsic biases between assessment modalities, inconsistent response classifications can be determined depending on the assessment modality used: mammogram (MG), magnetic resonance imaging (MRI), ultrasound (US), physical examination (PE), computed tomography (CT), blood biomarkers and others.

Accurately classifying tumors with distinct trajectories is necessary to identify biomarkers specific for one of these outcomes (9, 10). Further, assessing the tumor trajectory during the course of therapy can provide accurate real-time assessments to guide adaptive treatment strategies (4) (for current perspective see (11)). A more dynamical approach to measuring changes in tumor size could also capture the evolution of resistance during treatment.

The heterogeneity of solid tumors, such as breast cancer, can also obstruct the accurate measurement of tumor size when using a single imaging method alone. Physicians thus rely on a combination of imaging and physical examination modalities during the course of therapy to understand a tumor’s characteristics and to increase accuracy compared to using a single methodology (12, 13). The accuracy of each modality depends on patient specific factors, such as the level of inflammation or cancer subtype, and in clinical practice patients may be assessed by different modalities at different time points (14-16). However, when using a range of methodologies, it is challenging to combine information to a consensus reading or to integrate data sources across time points and methodologies.

For early-stage ER+ breast cancers, tumor diameter on clinical physical examination, ultrasound, mammogram, and MRI can be used to assess response to neoadjuvant therapy. However, the use of a single imaging modality less accurately reflects surgical pathologic response or long-term treatment efficacy (17-21). Among the various imaging modalities, MRI appears the most accurate for predicting pathologic response in breast cancer (21). However, MRI accuracy is still below 80% for predicting pathologic response, and MRI use in earlier stage breast cancer is variable across treatment centers (22). Further, the use of MRI in early-stage breast cancer remains controversial because of unclear effects on long term outcomes and concerns about over diagnosis of incidental indolent lesions (23, 24).

Here we provide a general approach to reconstruct the trajectories of patient tumors during treatment. The approach starts by generating a diversity of possible trajectories of tumor size over time using a Gaussian process model (25). To reconstruct the most likely tumor trajectory, we apply a Bayesian probabilistic framework to integrate all available measurements of tumor size during treatment and account for biases and differences in precision of each method (26). We test the power of the trajectory reconstruction approach to accurately recovers the underlying dynamics of tumor progression using *in silico* simulations. We then apply the model to reconstruct tumor trajectories using imaging data from a randomized clinical trial of ER+ breast cancer patients during neoadjuvant treatment with an aromatase inhibitor combined with a cell cycle inhibitor or placebo. From these data, we identify five distinct trajectories of tumor progression in this cohort, including a group of patients with a rapid rebound of disease after an initial response. The model also reveals the speed of growth changes during treatment, including shrinkage of tumor size in sensitive tumors and increased growth in resistant tumors. Applying this model to patients across treatments, we show that combination endocrine and cell cycle inhibitor therapy increases the frequency of resistance-related tumor trajectories compared to endocrine therapy alone. Further, high dose combination therapy increases the frequency of rebound disease outcomes. In summary, tumor trajectory reconstruction integrates multiple modalities of tumor measurement to describe the diversity of tumor response to therapy with increased resolution and dynamical precision.

## Methods

### Trial overview

Tumor diameter (mm) was monitored over six months during a multi-institutional phase 2 trial of women with early-stage ER+, HER2-breast cancer (27) which evaluated whether the addition of CDK inhibition to endocrine therapy in the neo-adjuvant setting promotes complete cell cycle arrest and improves the preoperative endocrine prognostic index (PEPI) (27) (28). Post-menopausal women with node positive or >2 cm ER+ and/or PR+, HER2 negative breast cancer (n=120) were randomized equally between three treatment arms, receiving: A) endocrine therapy alone (letrozole plus placebo), B) intermittent high dose combination therapy (letrozole plus ribociclib: 600 mg/day, three weeks on/ one off), or C) continuous lower dose combination therapy (letrozole plus ribociclib: 400 mg/d).

### Measurements of tumor size

To assess each patient’s tumor progression, a range of standard imaging and physical examination assessments were used at different time points throughout the trial. Large discrepancies were observed between the estimates of tumor size from each of these measurement approaches (Fig S1), motivating the application of our tumor trajectory reconstruction approach. As is normal in the clinical setting, each patient had a unique combination of magnetic resonance imaging (MRI), ultrasound (US), mammogram (MG), clinical physical examination (PE), and surgical pathology (SP) observations. Imaging and physical examination assessments were made repeatedly during the 180-day period of neo-adjuvant therapy (mean= 6.8 tumor measurements per patient). Of the 120 patients, 91% had sufficient clinical observations to reconstruct their tumor burden trajectory.

### Reconstructing tumor burden trajectories: Probabilistically combining tumor estimates from different measurement methods

To reconstruct patient’s tumor trajectories during treatment, we developed a dynamic response evaluation approach (Fig.1 and below). Tumor trajectories were reconstructed by combining all available sources of clinical imaging, physical examinations and pathological data, using a Gaussian Process Latent Variable Model (GPLVM) (29, 30). Potential tumor size trajectories were probabilistically evaluated, based on clinical data and known biases and differences in accuracy of different clinical measurement modalities, and the most likely tumor burden trajectories were learned using Bayesian statistical inference.

**Fig.1.**
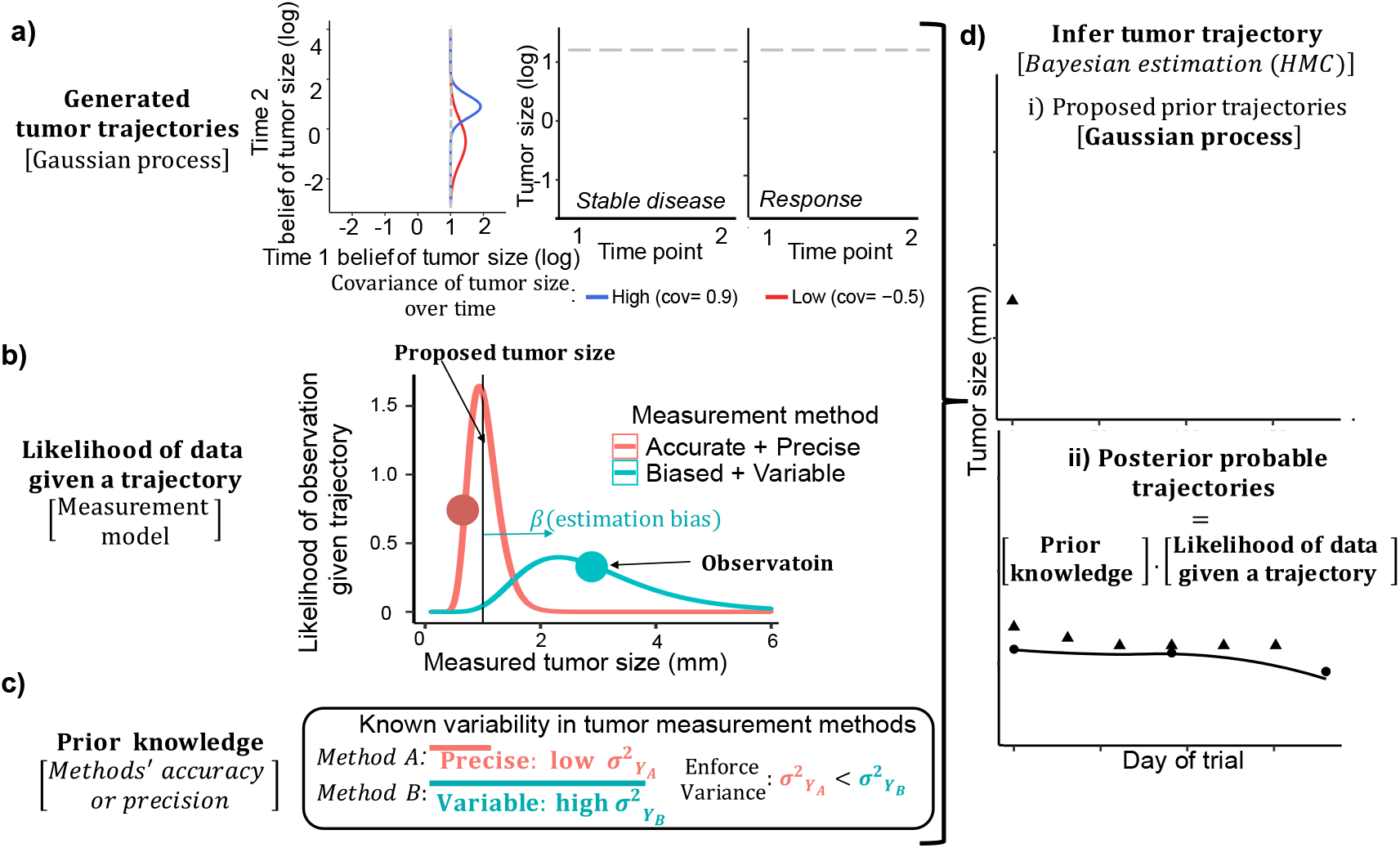
Overview of tumor reconstruction approach. **a)** The Gaussian Process model generates a wide range of potential tumor size trajectories by flexibly describing the correlation (covariance; left panel) between the tumor size at different time points. High covariance between time points (blue) generates stable tumor size during that timeframe (center panel), while low covariance (red) produces fluctuations in tumor size (right panel). **b)** Trajectories proposed by the GP are evaluated for consistency with clinical data. For each observation (point) made under each measurement method (color), the measurement model calculates the probability of the data, given the tumor is the size proposed by the Gaussian process. Combining these probabilities, information from multiple measurement methods is combined, allowing all available clinical data can be used to reconstruct tumor trajectories. Comparing observations across different measurement methods, the accuracy of specific methods is quantified (biases measured by *β*) as well as differences in precision (measurement noise measured by *σ*). **c)** Existing knowledge about the differing precision (*σ*) and bias (*β*; tumor size over/underestimation) of measurement techniques is incorporated into Bayesian priors put on the measurement model parameters. **d)** Tumor trajectories are learned using a Bayesian inference approach, by combining parts A-C. The Gaussian process proposes tumor trajectories (i) and the product of the prior knowledge and the likelihood of the clinical observations determines whether the trajectory should be accepted (ii). By iteratively proposing new and trajectories and accepting improvements, the model converges on the tumor size trajectories that are most likely to have occurred, as well as capturing our uncertainty in the trajectory (red lines=possible trajectories, black line =expectation). Biased clinical estimates (triangles) inform about the shape of the trajectory, whilst unbiased measurement modalities (circles) provide information to determine the size of the tumor more accurately at a given time.

#### 1) Generating proposal tumor trajectories

Potential tumor size trajectories were generated using a multidimensional Gaussian distribution 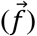 (Fig.1a). Each dimension (1: *n*) describes the log tumor size on an occasion (*i*) when the tumor was measured:

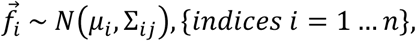

with *μ*_*i*_ reflecting the expected tumor size (log mean during treatment) and Σ_*ij*_ capturing the covariance of tumor size between occasions.

Smoothly varying tumor trajectories are generated when ordering the dimension indexes by time. The resulting Gaussian process (*f*(*t*)) is a probability distribution describing the potential state of the tumor over time (31). Possible tumor trajectories were generated by sampling nonlinear functions from this Gaussian process:

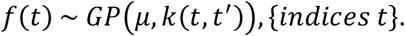

Here, *μ* scales the average tumor size and the covariance matrix *k*(*t, t*′) encodes how the tumor size changes between observation times. For example, a high correlation in tumor size between two time points yields little change in tumor size and produces a stable disease dynamic over this time frame (Fig.1a). The covariance structure was calculated using the squared exponential covariance function:

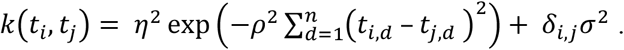

The length scale (*η*), controls the timescale at which the tumor burden fluctuates, whilst the signal variance (*ρ*) determines the amount of tumor size variation during treatment. Together these estimated parameters of the covariance function describe the smoothness of a tumor burden trajectory. Finally, a viable positive definite covariance matrix was ensured by adding *δ*_*i,j*_*σ*^2^ to the diagonal covariance elements (*δ*_*i,j*_= Kronecker delta function: 1 if *i* = *j*, else 0; *σ*^2^=1 × 10^−8^).

#### 2) Evaluating the likelihood of a tumor burden trajectory

To determine the probability of a proposed tumor trajectory, measurement models were used to describe the accuracy and precision of each tumor measurement method (*m*) (Fig.1b) (32). The measurement models then evaluate the likelihood of each tumor observation (*Y*_*P,m*_(*t*); method *m*, patient *P* at time *t*), given the tumor size followed the trajectory proposed by the Gaussian process (*f*(*t*)). Observations were described as lognormally distributed measurements of a patient’s true tumor size, with the addition of a censoring threshold of less than the minimal measurable size (*φ* = 0.1 *mm*):

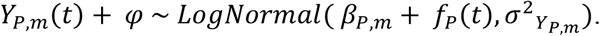

Each measurement methods’ bias (*β*_*P,m*_) and inaccuracy 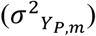 was estimated on a patient specific basis.

#### 3) Incorporating prior information about measurement precision and bias

Knowledge about the rank precision of different measurement methods was encoded into the measurement models (Fig.1c) using conclusions of four recently published comparative studies examining the agreement between surgical pathology measurements of tumor size and MRI, ultrasound, mammogram or clinical examinations (12, 14-16). The four key conclusions across these studies were that: i) MRI is the most accurate imaging method to estimate tumor burden, with little or no systematic bias compared to the observation made at pathology. ii) Ultrasound provides similar measurements, but potentially with greater bias and variability, iii) Mammograms results are significantly more variable and also potentially biased iv) Clinical physical examinations provide the least accurate estimates of tumor size, as they systematically underestimate tumor size.

Reflecting these conclusions, we constrained the variance parameter of more precise measurement modalities to be lower (*σ*^2^_*PE*_ > *σ*^2^_*MG*_ > *σ*^2^_*US*_ > *σ*^2^_*MRI*_ > *σ*^2^_*SP*_) and estimated bias in clinical assessment, mammogram and ultrasound measurements (*β*_*P,SP*_ & *β*_*P,MRI*_ = 0; *β*_*P, PE*_, *β*_*P,MG*_&*β*_*P,US*_ ≠ 0; sign of bias not constrained). Along with the biological requirement that tumor size changes gradually over time, we constrained the tumor size not to fluctuate at timescales shorter than one month.

##### Inferring tumor burden trajectories

The most probable tumor trajectories during the trial were identified using Bayesian inference to combine prior information about methodological biases with the likelihood assessment of trajectories made by the measurement model (Fig.1d) (33). The Gaussian process generated proposal trajectories, the measurement model quantifying the consistency of observations with the proposed trajectory, and current clinical knowledge of the accuracy and precision of measurement methods were all encoded in the Bayesian priors.

The fitted GPLVM tumor reconstruction provides patient specific estimates (with uncertainty) of: i) the tumor size throughout the trial (*f*(*t*)), the speed and extent of tumor size fluctuations (*η* and *ρ*) and the bias and precision of each measurement method. The confidence in the patients’ tumor trajectory is also captured in the Bayesian posterior distributions of the sampled Gaussian process. All parameters were inferred simultaneously using Hamiltonian Monte Carlo in Stan (34).

### Validating tumor trajectory reconstruction

To verify the reliability of the GPLVM tumor reconstructions, we generated hypothetical tumor trajectories using a theoretical model of tumor growth and subclonal evolution (35). We then simulated the process of measuring this *in silico* tumor using measurement methods with differing precision and accuracy. Finally, we assessed how well our dynamics response evaluation approach could reconstruct the trajectory of the simulated tumors, based on the measurement observations that were produced (Fig.2). We compared the GPLVM tumor reconstruction with RECIST response category and naive smoothing of tumor measurement data (using generalized additive models) (36).

**Fig.2.**
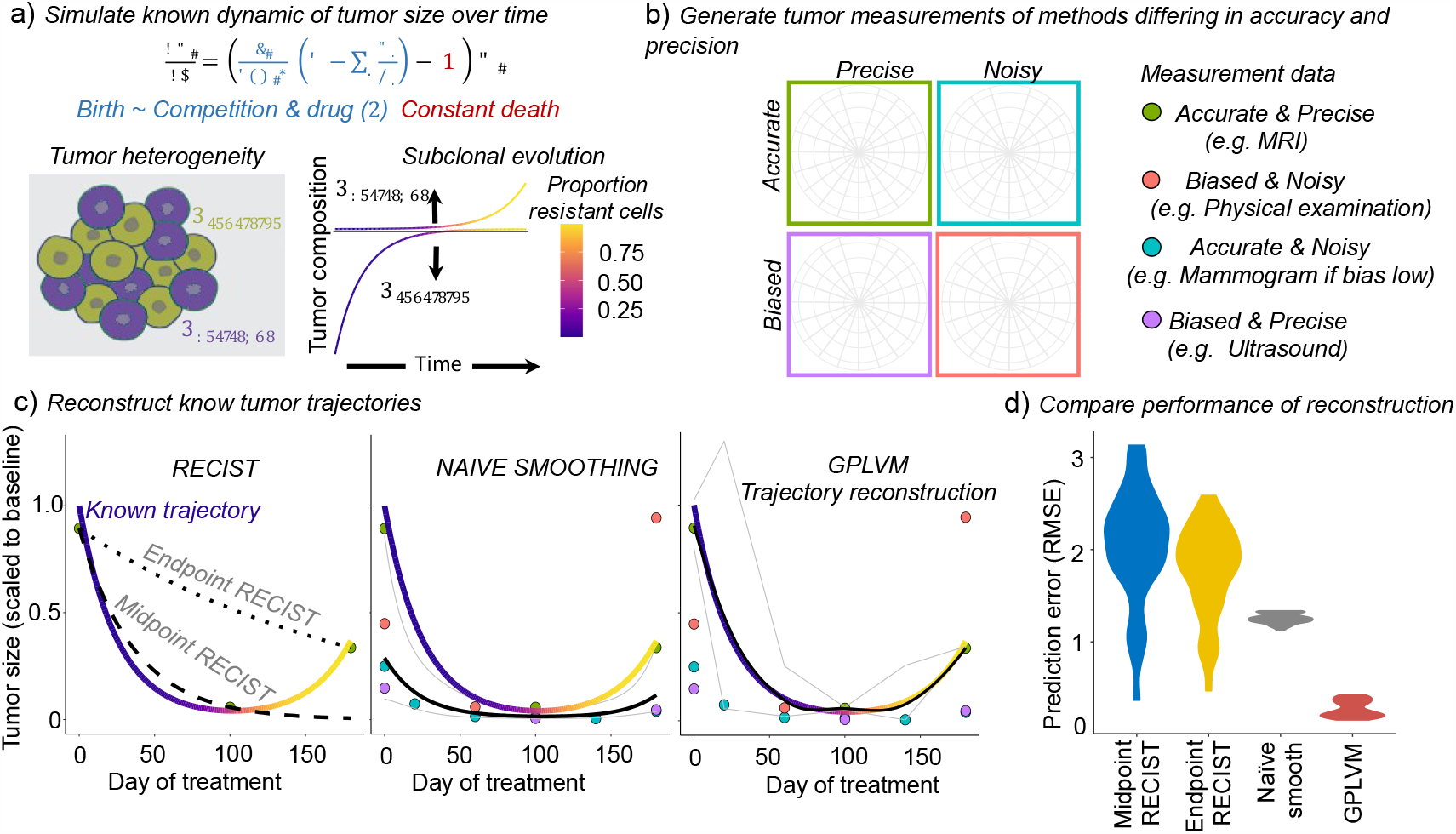
Validation of the performance of the tumor trajectory reconstruction approach, using simulated tumor trajectories and measurement observations to test the ability to recover known dynamics. **a)** Schematic and equation for the theoretical model of subclonal evolution of tumor resistance, used to generate *in silico* tumor trajectories. Subclonal tumor population (*i*) changes in size (*N*) following cell death and proliferation which depends of drug dose and the density of competing resistant and sensitive cells. Under drug selection, the tumor composition shifts from being dominated by sensitive (purple) to resistant (orange) subclones. Black line subdivides resistant and sensitive cells and distance to the colored line above and below indicates the abundance of resistant and sensitive cells respectively. Coloration of line signifies the proportion of resistant cells. **b)** Tumor observations are next generated by simulating the observation process for measurement methods with different levels of bias/accuracy and precision/noise. **c)** *In silico* tumor observations are used to reconstruct the known tumor trajectory (colored line signifies tumor resistance). Three methods to assess tumor trajectories are assessed: i) RECIST assessments (either comparing baseline with midpoint or endpoint tumor size), ii) naïve smoothing of the tumor observations from different measurement methods or iii) our GPLVM approach. Black lines indicate the predicted tumor trajectory and shaded regions indicate confidence intervals. RECIST assessment does not provide a measure of uncertainty, so we use dashed black and grey lines. Colored points indicate the data used by each approach. **d)** Comparison of the performance of the tumor reconstruction approaches as measured by the amount of prediction error between the known and predicted tumor trajectories (RMSE=residual mean square error of predictions). Violin plots show the distribution of prediction error for each approach, across 42 in silico tumor trajectory simulations varying the subclonal tumor composition and drug dose. Lower RMSE values indicate better performance.

We simulated tumor size trajectories over time, using a subclonal model of tumor growth and evolution: 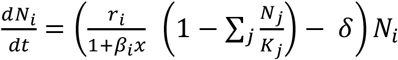. In this model we described the interaction between resistant and sensitive cells competing for limited resources. The tumor is initially dominated by sensitive cells, but a small subpopulation of resistant cells was allowed to pre-exist. Cell proliferation (**r**_**i**_) is reduced by the cell cycle inhibitory effects of treatment (**x**). Resistant cell proliferation is assumed to be less sensitive to the impacts of therapy (**β**_**R**_ = **β**_**S**_**/10**). For simplicity, similar death rates (***δ***) and competitive abilities 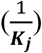 are assumed for the two cell types, as these assumptions do not influence conclusions of the simulation study.

We generated a range of different tumor trajectories by varying the initial percentage of resistant cells between 0.01 and 0.4% (n=6 resistance levels) and varying the drug dosage from 33% to 100% of the maximum dose that would cause shrinkage of a completely sensitive tumor within 50 days (n=7 dosage levels). A broad range of tumor trajectories were generated (n=42), based on simulations with each combination of drug dosage and initial resistance levels (Fig S2).

To assess the performance of the tumor reconstructions, we compared how well the known dynamics of tumor growth were recovered and compared this to trajectories predicted by: i) midpoint or end of treatment RECIST assessment and ii) a naive smoothing of all the clinical measurements using a generalized additive model. The performance of each approach was measured by the root mean-square-error (RMSE) between recovered trajectories and the known true tumor trajectory (RMSE closer to zero indicate less error in tumor trajectory reconstruction).

### Identifying distinct dynamic response classes in clinical trial patients

To compare tumor trajectories between patients, we standardized each trajectory, scaling by the tumor size at baseline. We utilized the patient’s tumor trajectory to quantify each tumor’s growth during the first phase (day 0-90) and second phase (day 90-180) of the trial. We calculated the tumor growth rate during each phase as well as the proportion of tumor remaining at end of treatment (relative to baseline). Based on these summary statistics, patients with similar overall tumor response trajectories were categorized into dynamic response classes, using a Gaussian mixture model (37).

The relative frequency of each dynamic response was calculated for tumors in each treatment arm. We examined whether certain dynamic responses where associated with a specific treatment regimen, using a chi-squared test. Pearson residuals were examined to identify which dynamic responses were strongly associated with a given treatment. All statistical analyses were conducted in R 3.5.1 (R Core Team 2018), using the RStan interface to perform Bayesian inference in Stan (34, 38). The code to implement the tumor reconstruction is provided along with the ER+ breast cancer patient clinical data (Supplemental data= Online Data Supplement 1; Supplemental code= Online Data Supplement 2).

## Results

### Tumor trajectory reconstruction and validation

To test that our approach allows the reconstruction of tumor shrinkage and/or growth during treatment, we used a theoretical model of the subclonal evolution to a resistant state to generate trajectories of *in silico* tumors during treatment (Fig.2a). Initially drug sensitive *in silico* tumors develop resistance, as the composition becomes dominated by the initially rare resistant subclone, following drug-induced evolutionary selection. We next generated observations representing serial measurements of the tumor using methods differing in accuracy (bias) and precision (noisiness) (Fig.2b). We then compared our ability to reconstruct the underlying tumor trajectories using the GPLV model against predictions made using naïve smoothing of the observation data or using RECIST assessments of tumor size change comparing baseline to either midpoint or endpoint measurements (Fig.2c). Using tumor observations taken throughout the trial allows the identification of the emergence of resistance and the rebound of tumor growth, something not possible using the RECIST assessment (Fig.2c left vs center and right panels). When a naïve smoothing approach was used, assuming all measurements were equally reliable, the general trend of the trajectory was recovered; however, the size of the tumor could be poorly measured due to the biases of frequently used measurement techniques (such as clinical physical examinations) (Fig.2c center vs right panel). Our approach allowed a description the smooth changes in tumor size over time, using the Gaussian process, and to correct for method specific biases using the method specific measurement models (Fig.2c right panel). This approach allowed a quantitatively accurate reconstruction of the tumor trajectory that captures: i) the initial rate of decline in tumor size upon initiation of therapy and ii) the timing and speed of tumor rebound growth following the emergence of resistance.

The approach was applied to reconstruct a broad range of tumor dynamics, generated by modelling tumors with differing initial frequencies of resistant subclones and by varying the drug dose (Fig.2d). Across all tumor reconstructions, the GPLVM approach more accurately recovered the underlying tumor trajectories, having 87% less prediction error compared to midpoint RECIST assessments, 85% less that endpoint RECIST assessments, and 78% less than the naive smoothing approach.

### Reconstructing tumor trajectories: Probabilistically combining tumor estimates from different measurement methods

Reconstruction of patient tumor trajectories provides a dynamic evaluation of response during treatment (Fig.3). Here we show the inferences that are obtained for each patient’s tumor by first presenting results of a representative tumor. All patients’ tumor reconstructions are provided in the supplement (SI Figure= Online Data Supplement 3). Figure 3a shows the tumor size estimated throughout treatment, including at time points in between observations and the overall average tumor size during the trial. The fitted model also measures the speed at which tumor size fluctuates and also the magnitude of those changes relative to the tumors initial size (Fig.3b). For this specific patient, tumor responded over a timeframe of around 70 days (lengthscale≈ 70), indicating a gradual reduction in tumor size rather than a rapid decline as may be expected under cytotoxic therapies. Furthermore, the signal variance measured the limited extent of tumor response during the trial, indicating that the reduction in tumor size was limited to only 16% of the baseline size (Signal variance≈ 0.15).

**Fig.3.**
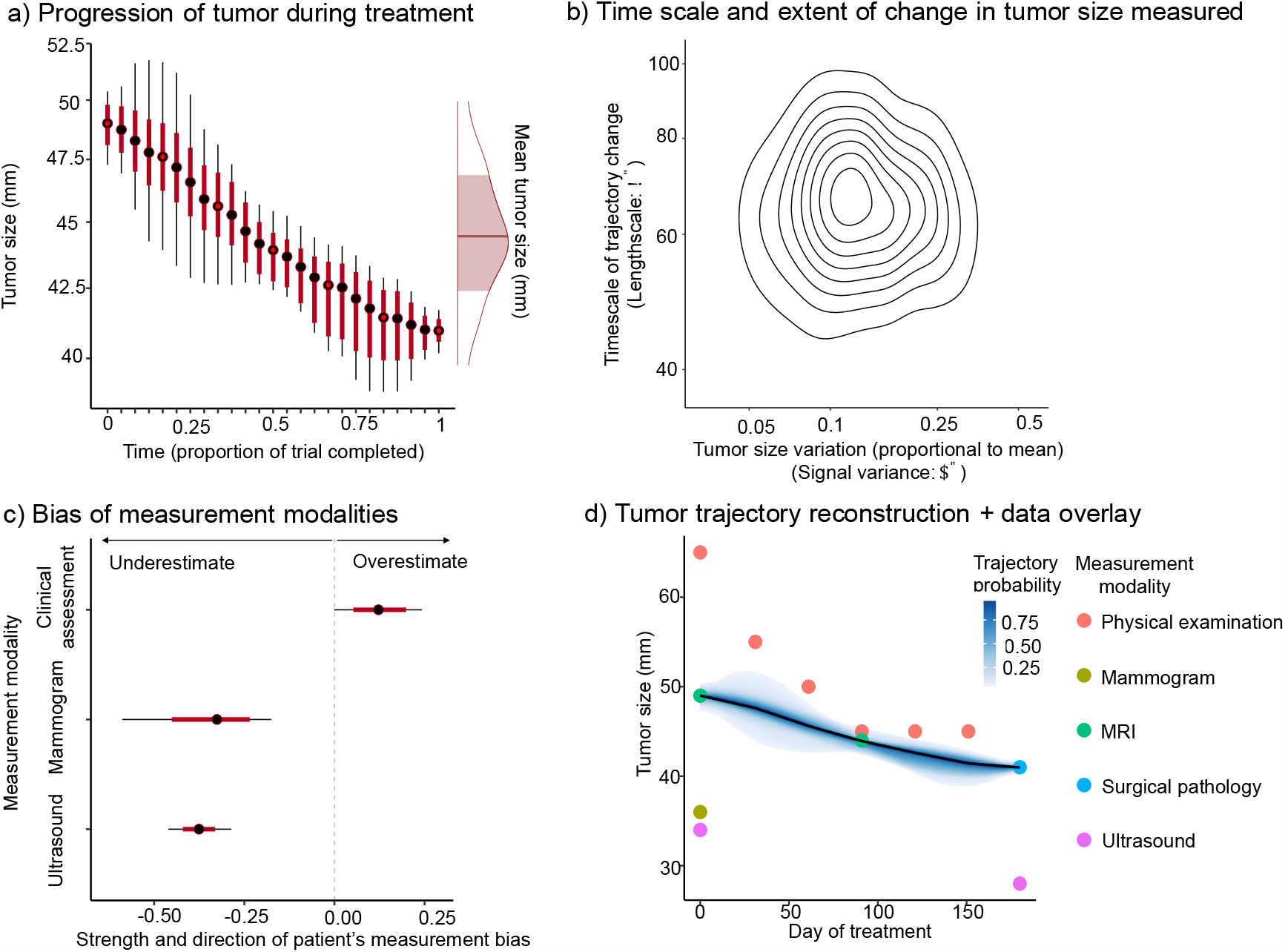
Inferences from tumor trajectory reconstruction applied to a representative patient. **a)** Throughout the trial, tumor size is estimated (points) and uncertainty measured (box and whiskers=95 and 99% posterior interval). Tumor size can be measured at time points when tumor is measured (red filled points), but can also be inferred at intermediate times between measurements (solid black points), due to the reconstruction of smooth tumor size transition. The average tumor size (with uncertainty:95% credible interval) is shown by the right hand density plot. **b)** Scatterplot of Bayesian estimates of the timeframe and extent of tumor response to therapy that are consistent with the clinical observations of that patient’s tumor. The speed change in tumor size is measured by the lengthscale parameter, indicating the timeframe over which the tumor response occurred and the Extent of tumor response to therapy is measured by the signal variance parameter, indicating the proportional change in tumor size relative to baseline. Each point indicates the timescale and extent of tumor change in a trajectory that is consistent with the data. The highest density of estimates indicates the most probable value of the parameters and the distribution of estimates measures uncertainty. Contour lines highlight the most probable regions (contours indicate 10% reductions in probability). **c)** Box and whisker plot showing the bias of tumor size estimates provided by different measurement methods. Negative values indicate underestimation of tumor size and positive values show overestimation (box and whiskers=95 and 99% posterior interval). Unbiased estimate indicated by dashed grey line. **d)** Reconstruction of the smooth tumor trajectory (black line) with uncertainty (shaded region= high probability density credible interval, shade indicates trajectory probability). Clinical measurements obtained by different measurement modalities are overlaid (colored points).

Measurements of the over/underestimation bias of each tumor measurement method were obtained, showing that for this patient, clinical physical examinations were overestimating tumor burden, whereas mammogram and ultrasound provided underestimates (Fig.3c). These biases can be visualized in Figure 3d where the clinical measurements are overlaid onto the reconstructed tumor trajectory. The surgical pathology measurement, which measures actual tumor size at time of surgical removal of the tumor, provides validation that the final tumor size was substantially higher than was estimated by ultrasound. Similarly, the baseline MRI tumor measurement was substantially larger than ultrasound and mammogram assessments, indicating an initial size of around 49 mm. As the model describes smooth tumor size transitions over time, we can reconstruct the most likely tumor size trajectory (solid black line), and the Bayesian inference approach allows assessment of the range of tumor trajectories that are consistent with the data, quantifying the extent of our uncertainty in tumor trajectories (shaded region=high probability credible interval).

### Insights into the diversity of tumor response trajectories

The trajectory of each patient’s tumor size during the trial was reconstructed, probabilistically combining information from all available sources of clinical imaging, physical examination and pathological data, which together captures the most probable tumor burden fluctuation over time. Inferred tumor size at end of trial closely mirrored pathological observations (Fig S3) and frequently corrected for the underestimation of tumor size that previous studies have revealed (16) (Fig S4) (Strong underestimation through physical examination in 60% of patient’s tumors).

Five distinct dynamic classes of tumor trajectories were identified (Fig.4a-b). These categories corresponded to: i) sustained shrinkage (continued decline during trial; final size ≤25% baseline), ii) partial shrinkage (initial velocity of decline slowed in second half of trial and final size between 30% and 75% baseline), iii) stable disease (minor tumor size change and final size >70% and <150% baseline), iv) rebound disease (initial decline to size <70% baseline and rapid tumor regrowth during the trial), and v) progressive disease (increasing tumor size throughout trial despite treatment; final size >150% baseline). Figure 4c shows that the tumor response categories are distinct by comparing the overall reduction in tumor size during the trial and the initial rate of tumor size decline in the first phase of treatment. Similarly, by examining the change in tumor size continually during treatment, the five categories of tumor response show different trajectories (Fig.4c). For example, patient tumors exhibiting rebound disease had significantly more rapid reductions in tumor size during the first 100 days of treatment than patient tumors exhibiting partial or sustained response (lower tumor growth rate in days 0-100: versus sustained response t=-2.159, p<0.05; versus partial response t=-3.921, p<0.001). However, the subsequent regrowth after around 120 days of treatment contrasts the slower but more durable decreases in tumor size observed in sustained response tumors throughout the trial. Patients classified as non-responders using an MRI RECIST 1.1 assessment (Baseline versus day 90) were distributed between the residual disease categories, but reassuringly none were classified as sustained responders.

**Fig.4:**
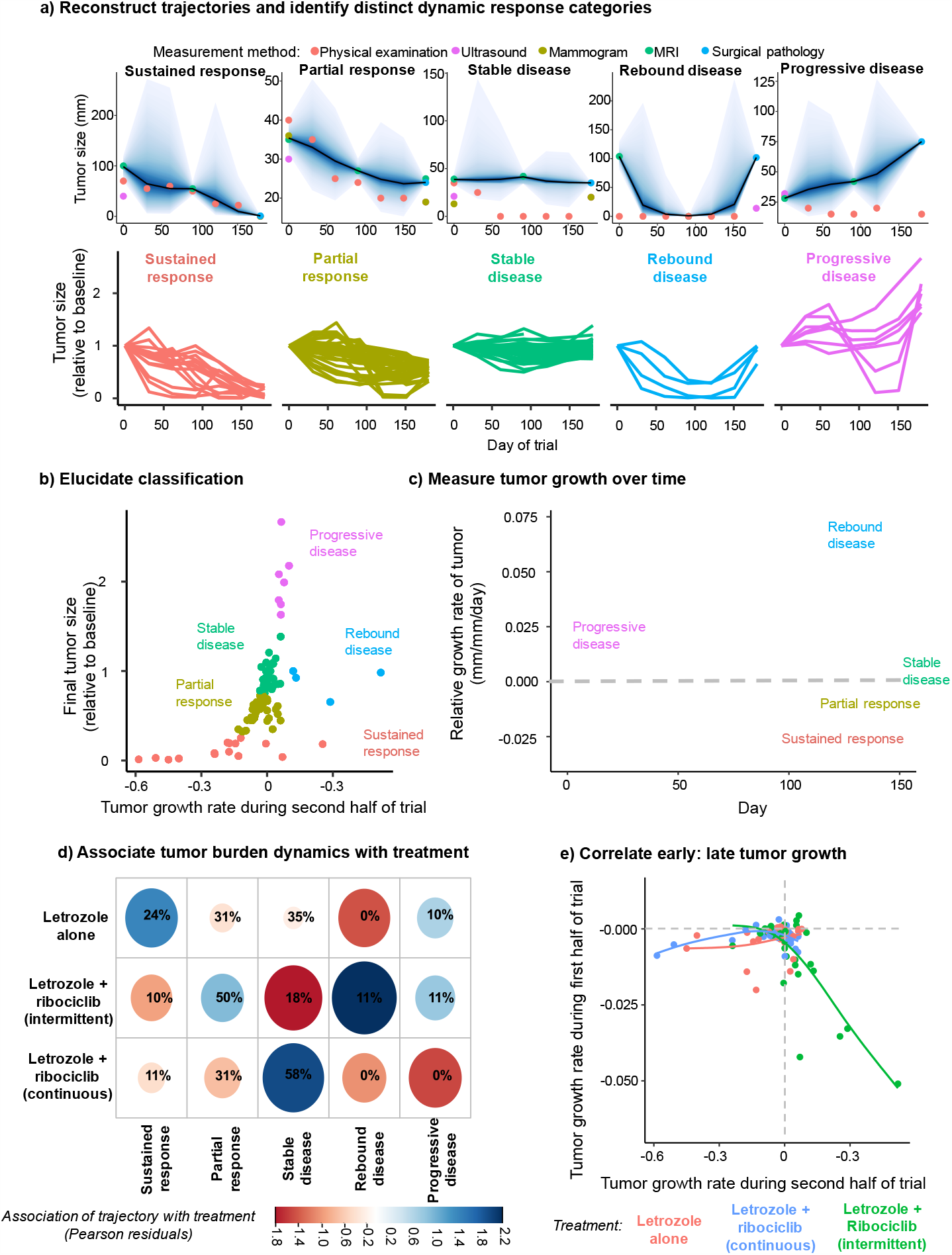
Distinct tumor trajectories associated with endocrine and combination endocrine and cell cycle inhibitor therapy. **a)** Five distinct response dynamics (columns) observed across clinical trial treatment arms: sustained response (continued decline to near complete response), partial response (initial response saturating, fraction tumor remaining > 40%), stable disease (little change during trial, tumor remaining>70%), rebound disease (initial response, subsequent regrowth) and progressive disease (continued growth despite treatment). Top panel: example tumors reconstructed and assigned to each category. Tumor trajectory reconstruction (solid black line) and uncertainty interval (shaded regions: high probability credible interval) are overlaid with the clinical data used to inform the trajectory (points: color indicates measurement method). Bottom panel: spiderplots of patient tumor size trajectories classified into each response category. **b)** Patient tumor trajectories summarized by measuring the reduction of tumor size (relative to baseline) and tumor decline rate in second half of trial (points=summarized patient tumor trajectory). Distinct tumor trajectory categories (colors) identified using a Gaussian mixture model. Confidence ellipsoids show tumor trajectory characteristics leading to each categorization. **c)** Comparison of the speed of tumor growth/decline (relative growth rate of the reconstructed tumor between observations) across the trial for tumors in each response category (color). Differences in timing and extent of initial tumor decline and extent of rebound growth when resistance emerges evident. Solid lines show average trends in tumor growth and shaded regions signify the heterogeneity of growth over time within a tumors response category (confidence intervals). Dashed grey line indicates no tumor growth (negative=shrinkage, positive=growing during time interval). **d)** Relative frequency of each trajectory (columns) in patient tumors from each treatment arm (rows). The size of the ellipse indicates the absolute value of the Pearson residuals, measuring how strongly a trajectory is associated with a given treatment arm. The color indicates whether the dynamic occurs more (blue) or less (red) frequently than expected in a given arm. **e)** Speed of growth of patients’ tumor (points) from each treatment arm (color) during first phase (day 0-90; y-axis) and second phase (day 90-180; x-axis) of the trial, as measured by the tumor reconstruction. Dashed horizontal and vertical lines indicate no net change in tumor size during the first and second phase of the trial respectively (positive values = tumor increased in size, negative values= shrinkage of tumor). Solid colored line shows the association between growth of the tumor during the first and second phase under each treatment (shaded region = 95% confidence intervals)

The frequency of patient tumors within each classification differed between the treatment arms (*χ*^2^=25.909, p<0.005) (Fig.4d), despite the average end of treatment tumor burden not differing between arms (39) (Fig S5). Therefore, tumor trajectory reconstruction reveals additional information about the time during which a treatment is effective and how rapidly resistance is emerging in a tumor. The use of combination therapy reduced the frequency of sustained response compared to endocrine therapy alone (z=3.040, p<0.005) and instead increased the frequency of stable disease trajectories under continuous low dose treatment (z=-3.221, p<0.005). The frequency of rebound disease trajectory was higher in the intermittent standard dose group than in the continuous low dose group (z=5.148, p<0.01). There were also more variable patient outcomes, as measured by an F-test of variance in final size (F=32.94, p<0.001; 2.7 × variability in final tumor size following intermittent vs continuous dosing) (Fig S5). Tumors exposed to intermittent high dose combination therapy decreased more rapidly during the first phase (day 0-90) of the trial, but this decrease was correlated with faster rebound growth in the second phase (day 90-180) (r=-0.76, F=16.87, p<0.0001) (Fig.4e). These added insights from the tumor reconstruction analysis suggests that combination therapy in this earlier stage ER+ breast cancer population, especially at higher doses, may potentially accelerate the evolution of endocrine resistance.

## Discussion

Reconstruction of tumor trajectories of early-stage ER+ HER2-breast cancers during neoadjuvant treatment provides a detailed assessment of the impacts of combining endocrine therapy with targeted cell cycle inhibition therapy. Although combination treatment had no significant impact on average tumor size by time of surgery (39), the assessment of tumor trajectories reveals that combination therapy produced more highly variable tumor responses with an increased frequency of rebound disease, especially under high dose intermittent treatment. Longer follow-up of the FELINE trial will inform if responses assessed by tumor trajectory reconstruction is prognostic. Further studies are needed to evaluate the role of drug dosage and timing on the evolution to a resistant tumor state.

Using reconstructed tumor trajectories to distinguish patients with these distinct evolutionary resistance backgrounds (innate vs acquired) is essential for effectively implementing system biology based targeted therapeutic strategies (40). Endpoint focused response assessments are unable to distinguish these diverse tumor trajectories or differentiate rebound disease reflecting acquired resistance from stable disease indicative of weak innate resistance. For patients exhibiting a rebound disease trajectory, continued treatment permits rapid tumor growth once resistance is acquired. Because the mechanisms of innate and acquired resistance may be different, the classification of tumors into groups with distinct trajectories, rather than clinical outcomes may enable development of treatment approaches that target innate and acquired resistance.

Another opportunity for improved patient care exists by using tumor trajectories to estimate the amount of bias for individual assessment modalities in individual patients. These estimates can be used when clinical circumstances require comparing one modality to a different modality at a previous time point. The weak correlation of tumor size estimates from surgical pathology with estimates from physical examination, and to a lesser extent ultrasound and mammograms, shows that single modality-based approaches are limited in accuracy. This reinforces the need to synthesize all available tumor measurements carefully, to account for the discrepancy in measurement accuracy across modalities.

Knowledge of the tumor trajectory, correcting for these biases, can guide adaptive therapy strategies, which work by initiating and ceasing therapy when the tumor reaches a critical size (11). As new data become available allowing the tracking of patient progress, the model can estimate how likely it is that the threshold size will be passed before the next assessment, allowing the application of adaptive therapies as threshold sizes are passed. Furthermore, there is no limitation to the number of measurement data types that can be used to inform the tumor trajectories. Other frequently monitored peripheral blood biomarkers of tumor burden can easily be integrated into the tumor reconstruction framework if sufficiently reliable markers are available.

These added insights from the tumor reconstruction analysis suggests that combination endocrine therapy and cell cycle inhibitors in this early-stage ER+ breast cancer population, especially at higher doses of the cell cycle inhibitor, may potentially accelerate the evolution of endocrine resistance. Interestingly, in the adjuvant treatment of early stage breast cancer, the MonarchE trial of abemaciclib, which is continuously dosed, showed an improvement in invasive disease free survival (41) while the PALLAS trial of palbiciclib, which is intermittently dosed like ribociclib, did not (42). Further research is needed determine whether the different results of these studies is related to the differences in dosing. Our results might inform the design of future neoadjuvant and adjuvant clinical trials by adopting continuous rather than intermittent dosing of the CDK4/6 inhibitors in combination with endocrine therapy. Furthermore, in early stage ER+ breast cancer, biological response to a neoadjuvant therapy is more prognostic than initial presentation factors (43). Hence, the adoption of a neoadjuvant strategy to help define tumor trajectories may add prognostic and predictive information to the ones currently available and allow us to change the treatment accordingly.

Reconstruction of patient tumor trajectories allows assessment of response throughout treatment based on multiple different assessment modalities instead of relying on just the start and endpoint measurements. This dynamic approach enables refined description of the diversity of tumor response to therapy and enables identification of personalized tumor trajectories not usually captured such as rebound disease dynamics. All available observations of tumor size progression can be combined with existing knowledge of method specific biases and precision to recover the most probable trajectories and to quantify uncertainty. Our tumor reconstruction approach could assist with treatment decisions based on interim imaging changes in neoadjuvant trials using adaptive therapy approaches.

## Notes

### Disclaimers

Ruth O’Regan participates on the advisory board for Cyclacel, PUMA, Biotheranostics, Lilly, Pfizer, Genentech, Novartis; Priyanka Sharma declares research funding from Novartis, Merck, Bristol Myers Squibb. Priyanka Sharma consults for Seattle Genetics, Merck, Novartis, AstraZeneca, Immunomedics, Exact Biosciences. Laura Spring participates on the advisory board for Novartis, Lumicell, Puma Biotechnology and Avrobio. Kari Wisinski participated on an advisory board for Eisai, Pfizer and Astra Zeneca. Kevin Kalinsky is a medical advisor to Immunomedics, Pfizer, Novartis, Eisai, Eli-Lilly, Amgen, Merck, Seattle Genetics and Astra Zeneca; his spouse is employed by Grail and previously by Array Biopharma and Pfizer. Anne O’Dea Consults for the Pfizer, PUMA Biotechnology, Astra Zeneca, and Daiichi Sankyo. All other authors have no disclaimers to report.

### Any prior presentations

This work has not previously been presented at scientific meetings or published

## Acknowledgements

We thank the anonymous patients from the trial that made this study possible.

## Author contributions

J.I.G. developed the tumor reconstruction approach, performed mathematical modeling and simulation studies and conducted patient analyses. J.I.G., F.R.A., A.L.C., A.H.B., contributed to study design, processed data and wrote the manuscript. A.L.C. contributed to data analysis and study design, provided clinical insight, and contributed to writing the manuscript. Q.K., conceived and coordinated the clinical trial, supervised contributed clinical support and infrastructure and provided clinical tumor measurement data, as well as contributed to writing the manuscript. OW was responsible for imaging assessment and contributed to manuscript editing. AOD, PS, MT, KK, KW, RO, IM, LS, AB, YY and MJ contributed patient data for the analysis and contributed to manuscript editing.

## Data availability

The data underlying this article are available in the article and in its online supplementary material. Clinical time course measurement are provided in Supplemental data= Online Data Supplement 1. Code to implement tumor reconstruction and to run the methodological validation simulations is also included in Supplemental code= Online Data Supplement 2.

## Supporting information

Despite clinical physical examinations suggesting complete response in 39% of patients, all examined patients were found to have some residual disease at the end of therapy surgery. MRI assessments could only be collected infrequently, but showed close agreement to the pathology results, with only 4% of patients being predicted to have experienced complete response. The inferred tumor burden trajectories, obtained from our mathematical model, show a more complete picture. The model predicts that none of the patients experience complete response, with the estimated tumor burden at the end of treatment correlating strongly with pathological results (Fig S1). Rather than obtaining just a classification of response at end of therapy, the model shows the most probably course of tumor size across the trial period, allowing a distinction to be made between patients that showed no response initially and those that experienced an initial response followed by relapse before the end of the trial.

**Fig S1:**
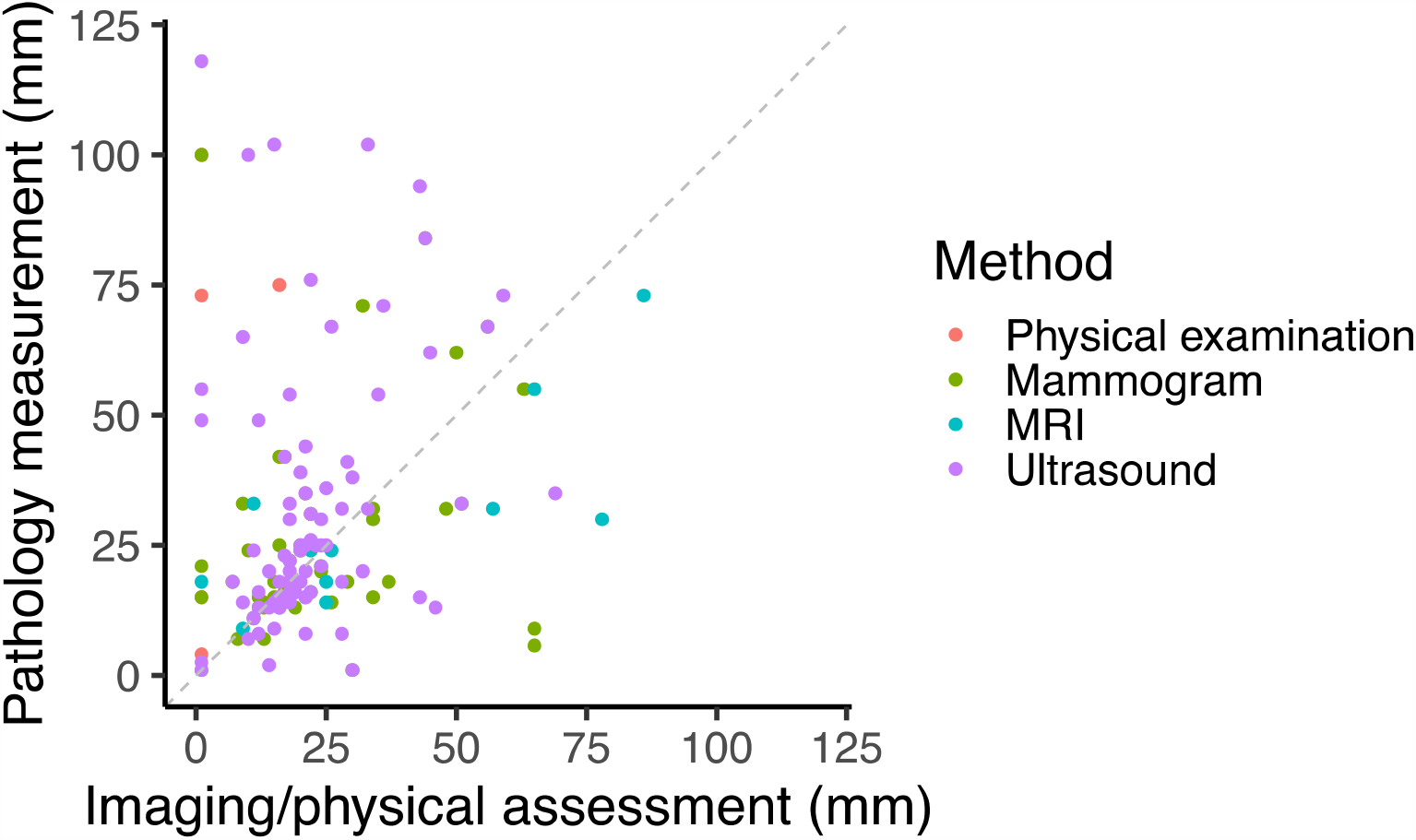
Large discrepancy (weak correlation) between surgical pathology observations of tumor size (mm) and measurements obtained by different imaging and clinical physical examination (colored points) just prior to surgery. Perfect agreement pathology observations and imaging/physical examination is indicated by the diagonal grey dashed line.

**Fig S2:**
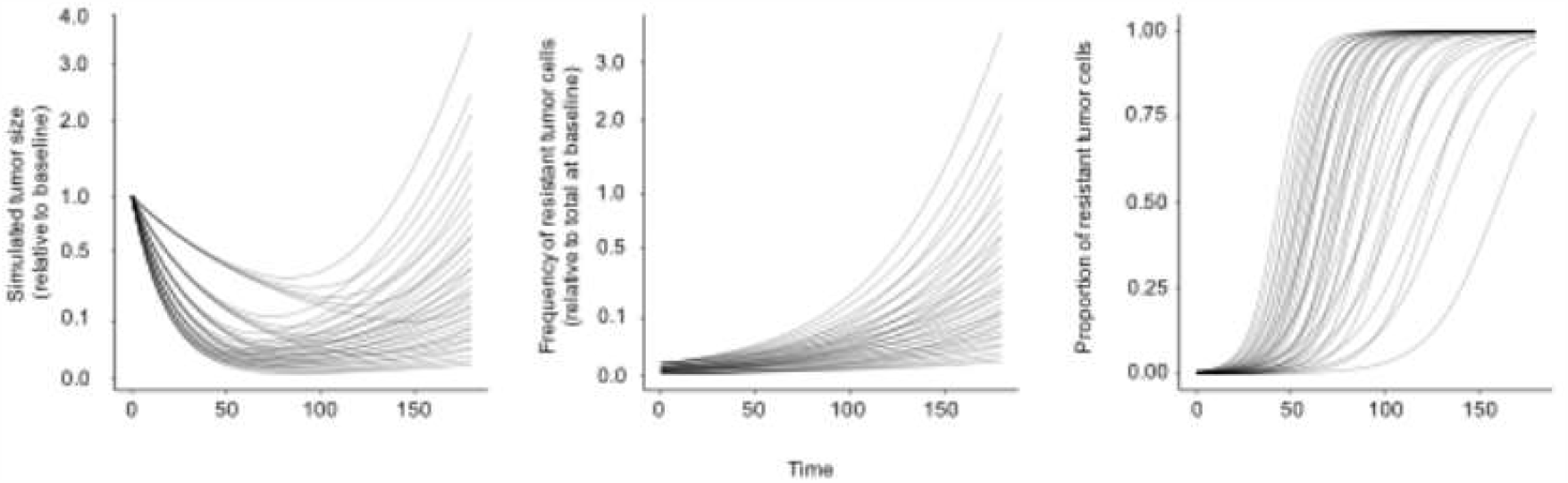
Diverse trajectories of in silico tumors generated by simulating the theoretical model of subclonal evolution of tumor growth. Left panel shows the diversity of changes in tumor size during treatment across simulations. Middle panel shows the emergence of a resistant subclone over time. The population size of the resistant subclone increases at different speeds, depending on the dose of therapy that drives selection. The initial frequency of this population also varies between simulations. Right panel shows the fraction of the total tumor that is comprised of resistant cells during the timecourse of therapy, showing the variation in the rate at which the resistant subclone becomes dominant.

**Fig S3:**
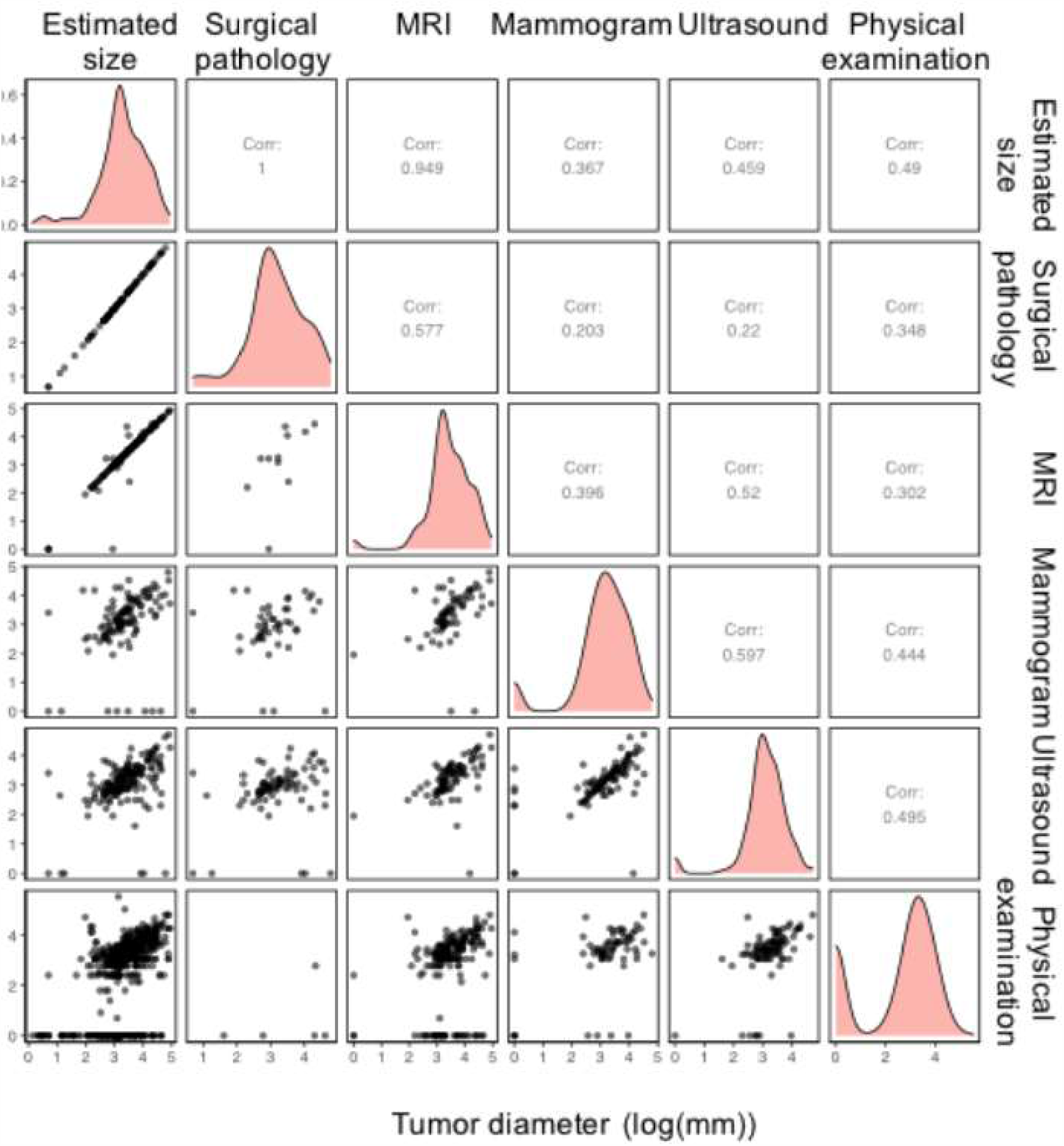
Strong agreement between tumor reconstructions and surgical and imaging measurements. Correlation of tumor size estimates from tumor trajectory reconstruction (first column/row) and surgical pathology observations (second column) and measurements from other imaging and physical examination assessments. Pairwise scatterplots show the agreement between each combination of methodologies (lower triangle of subplots). The correlation between these pairs of measurements is shown in the mirroring upper triangle of subplots. The overall distribution of tumor size estimates across patient tumors is shown by the density plots in the diagonal subplots.

**Fig S4:**
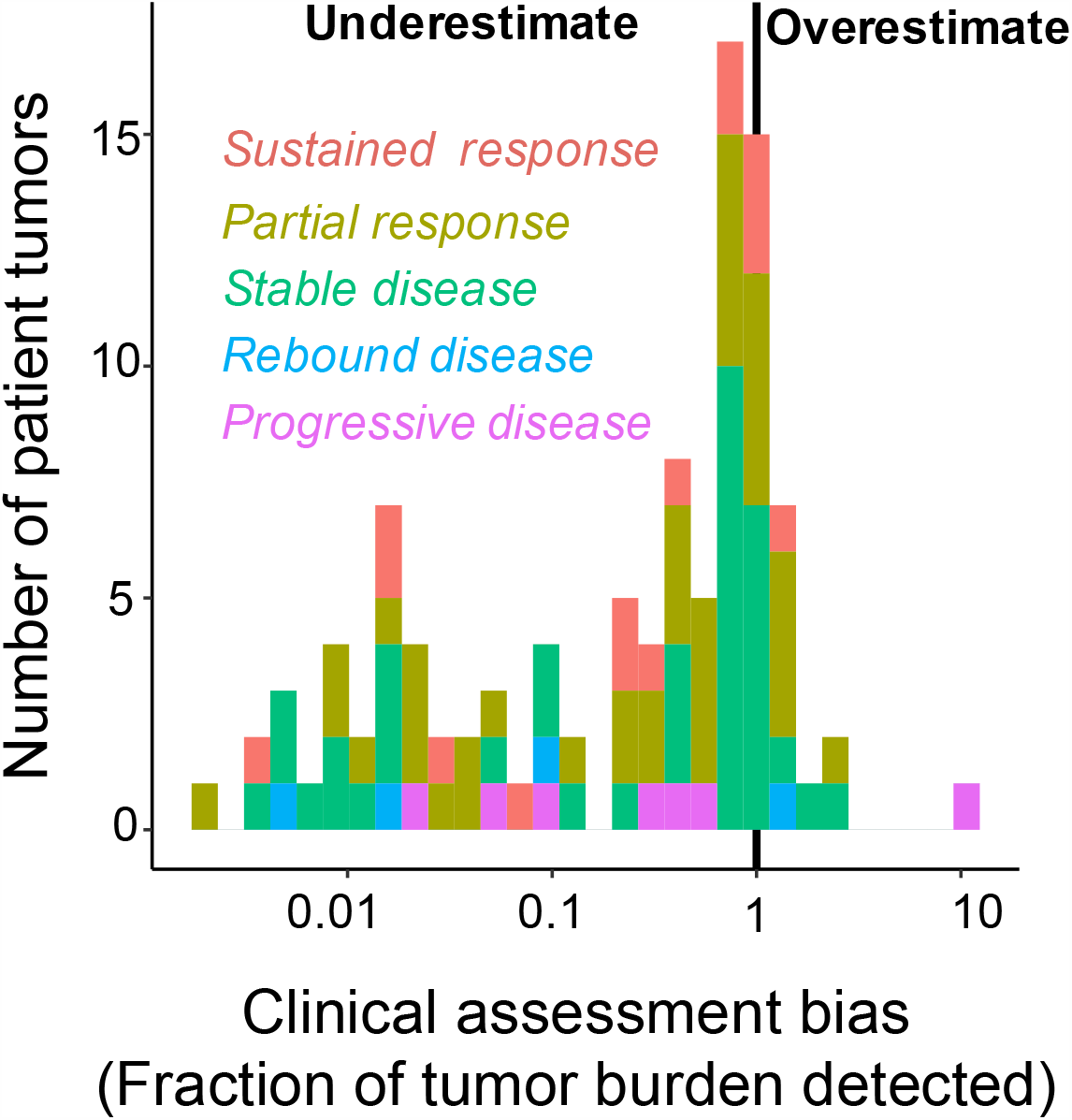
The tumor size measured by physical examination was consistently an underestimate of tumor size, independent of the tumor trajectory during therapy. Histogram shows the estimated fraction the tumor measured for patients with tumors exhibiting each tumor trajectory (color). Biases are estimated individually for each patients’ tumor and no constraint is added to enforce that physical examination underestimates tumor size. The bimodal distribution of measurement bias indicates that physical examinations were unbiased for around 60% of the patients of the study but provided large underestimates of the tumor size in the remaining 40%.

**Fig S5:**
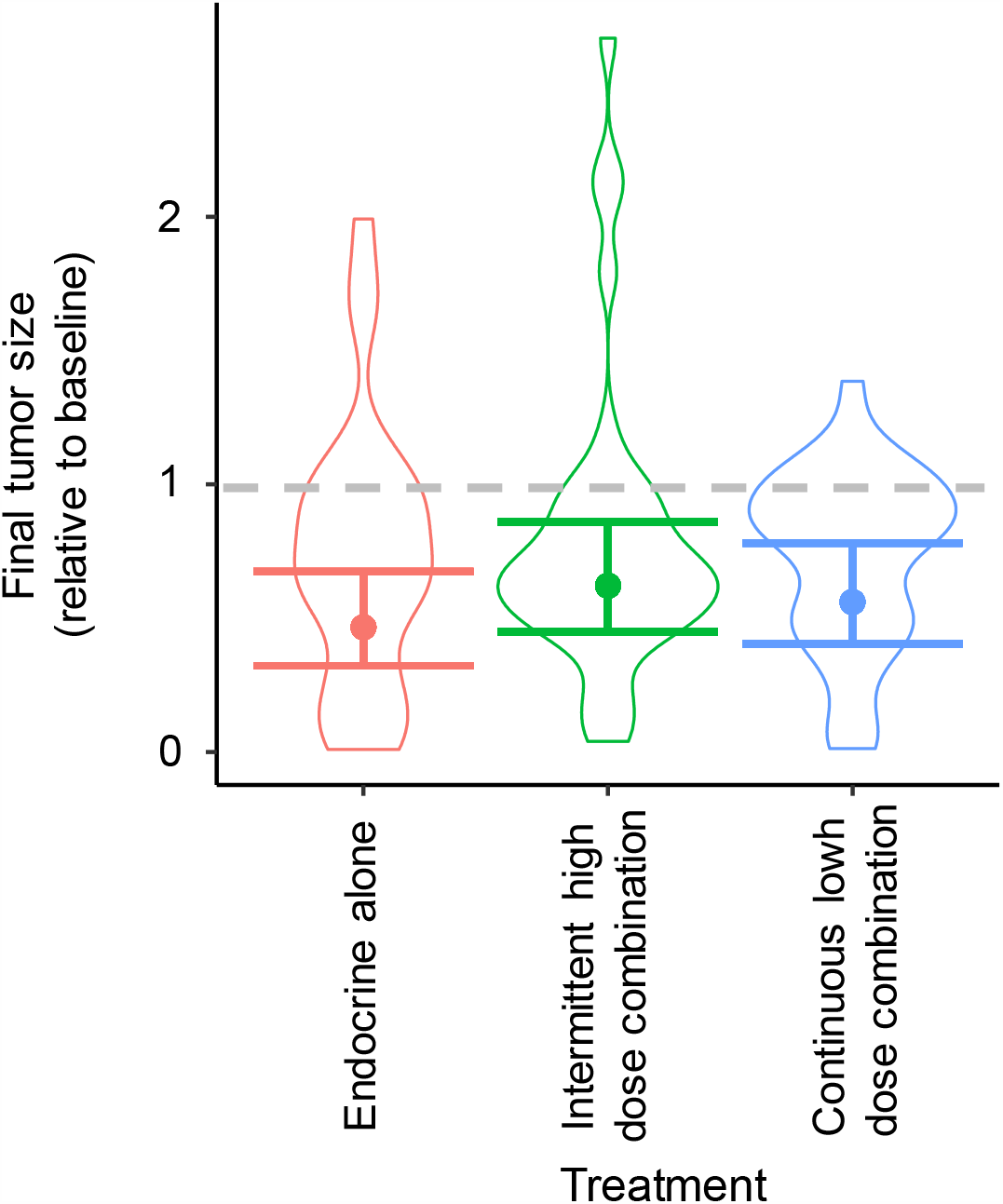
Combination therapy results in equal average reductions in tumor size compared to endocrine therapy alone. Violin curves show the distribution of final sizes of patients’ tumors (relative to baseline) in each treatment arm (color). Log linear regression used to compare average final tumor size (points show the expected final tumor size for each treatment. Overlapping confidence interval error bars shows that the final size did not significantly differ between groups.

